# Protein Abundance Inference via Expectation Maximization in Fluorosequencing

**DOI:** 10.1101/2025.07.10.664057

**Authors:** Javier Kipen, Matthew Beauregard Smith, Thomas Blom, Sophia Bailing Zhou, Edward M. Marcotte, Joakim Jaldén

## Abstract

1

Fluorosequencing produces millions of single-peptide reads, yet a principled strategy for converting these data into quantitative protein abundances has been lacking. We introduce a probabilistic framework that adapts expectation maximization to the fluorosequencing measurement process, estimating relative protein abundances with peptide inference results delivered by previously developed peptide-classification tools. The algorithm iteratively updates protein abundances, maximising the likelihood of the observed reads by obtaining more accurate protein abundance estimations.

We first assess performance on simulated five-protein mixtures that reflect realistic labelling and system errors. A simple Python implementation processes one million reads in under ten seconds on a standard workstation and lowers the mean absolute error in relative abundance by more than an order of magnitude compared with a uniform-abundance guess, demonstrating robustness in protein inference for small-scale settings.

Scalability is then evaluated with simulations of the complete human proteome (20 642 proteins). Ten million reads are processed in less than four hours on a NVIDIA DGX system using one Tesla V100 GPU, confirming that the method remains tractable at proteome scale. Using error rates characteristic of current fluorosequencing, the algorithm produces marginal improvements in relative abundance accuracy. However, when error rates were artificially lowered, estimation error decreased significantly. This result suggests that improvements in fluorosequencing chemistry could directly translate into substantially more accurate quantitative proteomics with this computational framework.

Together, these results establish EM-based inference as a scalable model-driven bridge between peptide-level classification and protein-level quantification in fluorosequencing, laying computational groundwork for high-throughput single-molecule proteomics. Furthermore, the proposed protein inference framework can also be used as a refinement step within other inference methods, enhancing their protein abundance estimates.

**Author summary:** Proteins carry out many of the functions inside our cells, but measuring how many copies of each protein are present remains a difficult task. The protein sequencing technology called fluorosequencing observes individual protein fragments one by one, producing millions of short “snapshots” in a single experiment. While this flood of data holds great promise, it is not yet clear how to turn the snapshots into reliable counts of the original proteins. In this study we developed a mathematical procedure, based on a classic “guess-and-improve” strategy called expectation maximization, that bridges this gap. Our program takes the computer’s best guesses for each fragment and repeatedly refines protein abundances until they best explain all of the measurements. Using realistic computer-generated data, we show that the method is both fast and accurate: it finishes in seconds for small mixtures and in only a few hours for the entire human set of proteins, both with accessible hardware. Because our approach can work with any future improvements in fragment interpretation, it lays the essential groundwork for bringing single-molecule protein sequencing into routine biological and medical use.

## 3 Introduction

Advances in next-generation sequencing technologies have rapidly transformed genetic analysis, enabling the complete sequencing of even the most complex genomes in a single day and drastically reducing the cost, time, and effort compared to first-generation methods [1]. In contrast, protein sequencing technologies have lagged behind, despite proteins being the primary functional molecules in cells and often more directly indicative of biological state. Many biological and medical applications, from cancer biomarker discovery to the analysis of complex signaling pathways, rely not only on identifying proteins but also on quantifying their abundances with high precision. Existing approaches, such as mass spectrometry and affinity-based assays, have made considerable progress, yet they often struggle to simultaneously achieve high sensitivity, digital quantification, and multiplexed measurement within a single platform [2].

Single-molecule protein sequencing (SMPS) has recently emerged as a promising class of technologies aimed at bridging this divide [3, 4]. Inspired by next-generation DNA sequencing, these techniques aim to identify and quantify individual protein molecules directly, without amplification, and at scale. While recent advances in nanopore-based methods [5, 6, 7] and emerging commercial platforms [8] hold significant promise, fluorosequencing [9, 10, 11] offers a compelling SMPS alternative. This last method combines fluorescent labeling of amino acids, Edman degradation, and imaging-based detection to extract partial sequence information from individual peptide molecules.

Some of the recent progress in fluorosequencing include estimations of experimental error rates [12], analysis and reduction of dye-dye interactions [13] and algorithms for classifying peptides from fluorescence readouts [14, 15]. The latter introduce peptide inference methods that account for model-based experimental failure rates. These methods are used as algorithmic components in this paper.

Although protein inference has been extensively studied in the context of mass spectrometry, with numerous methods developed for mapping peptides to proteins [16, 17], analogous approaches for fluorosequencing remain underexplored. In this work, we adapt expectation maximization (EM), a well-established probabilistic inference technique, to the fluorosequencing setting. We present a framework that estimates protein abundances by leveraging posterior probabilities produced by peptide-level classifiers such as Whatprot [14] or Probeam [15], and iteratively refines these estimates to maximize the likelihood of the observed fluorescence data. The EM algorithm provides a principled, computationally efficient, and scalable solution that aligns well with the structure and throughput potential of fluorosequencing technologies.

We validate our framework using simulated datasets and demonstrate that it robustly recovers protein abundances under realistic error conditions in small-scale protein mixtures. In addition, we present results from proteome-wide simulations, which validate the scalability of our method. Although improvements in proteome-wide abundance estimates are modest under present fluorosequencing error rates, our simulations demonstrate that reductions in these errors lead to markedly more accurate protein-level inference. While these results are encouraging, further methodological advances will be necessary to fully meet the demands of large-scale proteomic analysis in practical applications.

## 4 Results

### 4.1 Datasets

Simulated datasets were generated using the Whatprot framework [14] to assess our algorithm’s performance. We used simulations to verify our method since real proteome-scale experimental data is not yet available, and simulations provide the necessary ground-truth protein distributions for comparison, which would be extremely challenging to obtain under real-world conditions.

Ten distinct datasets were created for both experiments with five proteins and the whole proteome, each with different relative protein abundances, i.e., distributions. These true protein probability distributions were generated by sampling *N*_P_ exponential random variables and normalizing them by their sum, ensuring valid probability distributions while allowing for relative abundance variations across several orders of magnitude. For the whole-proteome case, this approach typically results in the most abundant protein being approximately 10^6^ times more abundant than the least abundant for each dataset.

A shared dataset of a thousand and a hundred reads per experimental fluorescence string was generated for the five protein and whole proteome experiments, respectively. Reads for each of the ten datasets were obtained by sampling the shared dataset with their respective fluorescence string distribution, given by Equation 6. The error metric used to assess performance, mean absolute error (MAE), is defined in Equation 20. All plots which illustrate the MAE consist of data points representing the mean error across datasets, while the error bands indicate the standard deviation. Lower MAE corresponds to a more accurate estimation of the protein abundance.

Throughout the Results section, we refer to the output of peptide-level classifiers such as Whatprot and Probeam as peptide inference. While this output is formally defined in the Methods section as fluorescence string classification, we adopt the term peptide inference here for clarity and to remain consistent with terminology commonly used in related literature.

We used Probeam to infer the posterior probabilities of peptides for each read, as described in Equation 12. This choice was primarily motivated by the flexibility of Probeam’s in-house implementation, which allowed us to modify the code to output the top *N*_b_ peptides rather than only the most likely one. Although prior work suggests that Probeam performs slightly worse than Whatprot for the peptide-inference task, we show in Appendix 9.4 that both classifiers yield comparable accuracy on the datasets used in this study. Based on this observation and the ease of integration, we proceed with Probeam as the sole peptide inference method throughout our experiments.

GPU acceleration was used to optimize computation time in the whole proteome experiment, and all algorithm calculations were performed on an NVIDIA DGX station with Tesla V100 GPUs. Appendix 9.8 provides further details on the GPU-optimized implementation. The five-protein inference was carried out on the same system but implemented in a Python script without any GPU acceleration, in order to provide a simpler and more easily reproducible codebase for small-scale scenarios.

### 4.2 Five proteins abundance estimation

While the fluorosequencing platform is ultimately intended for whole-proteome abundance estimation, it is also well-suited for early-stage development and use cases involving smaller sets of proteins. To explore this regime, we simulate a simplified scenario comprising only five proteins (PSME4, HBA2, HBB, PSME3, and PSMB10), as might arise in the case of a mixture of isolated proteins or a highly-enriched, partially purified proteomics sample.

In this experiment, we employ two peptide inference strategies to compute the posterior estimates needed for expectation maximization (EM): an oracle and a modified version of Probeam, both explained in section

6.7. The error rates of the simulations were parameters used in previous publications and are specified in the Appendix 9.3.1.

#### 4.2.1 Comparison of abundance estimation

A direct comparison of convergence curves across all methods is presented in Figure 1a, which includes the oracle, full Probeam, and sparse Probeam (*N*_b_ = 40). The presented computing time on the legend takes into consideration the whole ten datasets. While both Probeam variants converge similarly and lag behind the oracle in terms of final accuracy, the sparse version requires less computations and memory. For five proteins the differences between the sparse and non-sparse are not significant, but the sparse method is more scalable in terms of the amount of proteins in the dataset.

**Figure 1.**
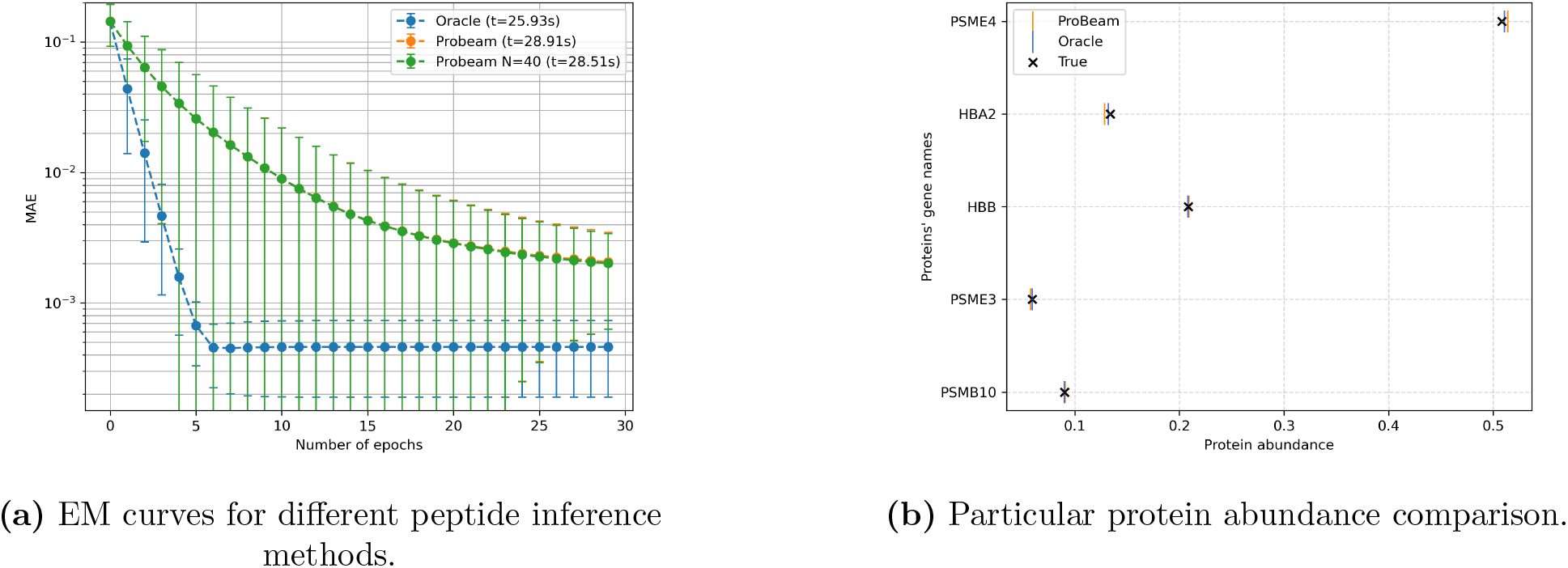
Performance comparison across different posterior estimates for a 5-protein experiment.

Finally, Figure 1b illustrates the ground truth versus the estimated protein abundances for each method in one of the ten data sets. All estimates closely align with the true values, confirming that even approximate posteriors derived from practical peptide inferences with currently achievable experimental errors can provide useful and reliable abundance estimates for datasets with few proteins.

In summary, this section demonstrates that the proposed EM-based inference method performs robustly in small-scale settings. For applications involving few proteins -whether for prototyping, targeted studies, or constrained biological contexts-this approach offers a practical and effective solution for protein abundance estimation.

#### 4.2.2 Oracle performance

We analyze how an imperfect peptide classifier oracle with error error rate *e* affects the protein inference quality. As shown in Figure 2a, increasing the error rate leads to a marked degradation in the accuracy of the protein abundance estimates. This behavior is expected: inaccurate posterior probabilities of peptides result in larger deviations to the true protein abundance. Additionally, the convergence curves do not always decrease monotonically. This is not contradictory to EM theory, which guarantees monotonically increasing data likelihood, not necessarily monotonic improvement in an external metric such as the mean absolute error (MAE). The data likelihood is computationally expensive to evaluate, so we instead report the MAE, which provides an intuitive but not necessarily monotone trend.

**Figure 2.**
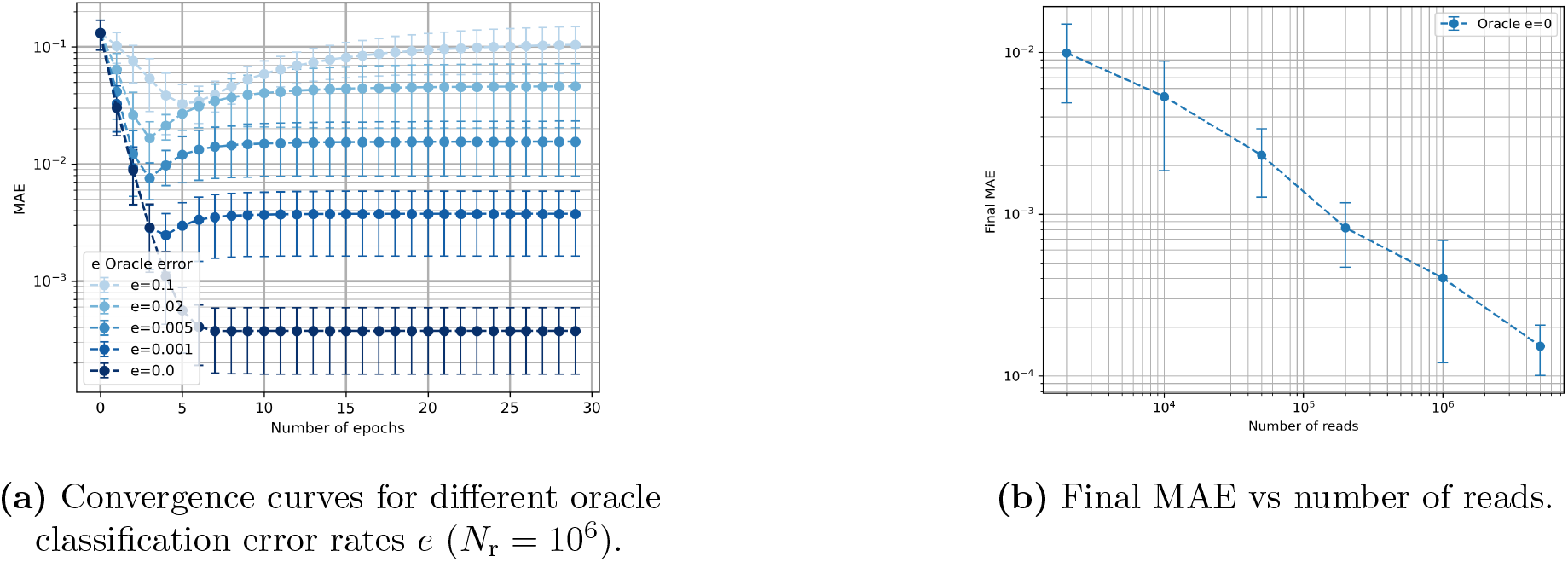
Sensitivity to oracle error rate and number of reads in a 5-protein experiment classified with an oracle.

Next, we evaluate the impact of the number of reads on estimation accuracy, as shown in Figure 2b. With a perfect oracle (*e* = 0), increasing the number of reads consistently enhances the estimation performance. This behavior shows that for this dataset a bigger dataset corresponds to improved abundance accuracy when the peptides are perfectly distinguishable.

#### 4.2.3 Sparsity sensitivity

We also examine inference using posterior probabilities derived from Probeam, focusing on the role of the sparsity of peptide observations in the posterior estimates. Figure 3 shows the convergence behavior and final MAE, for different sparsity levels. In sparse approximations, low-likelihood peptides are omitted, redistributing their residual probability across all peptides. This leads to a similar degradation in accuracy as observed with oracle error, when the residual probability is considerable. Specifically, aggressive sparsification (i.e., very low *N*_b_) results in significantly higher MAE as shown in Figure 3b, while moderate sparsity levels (*N*_b_ *≥* 40) yield estimates nearly indistinguishable from the full (non-sparse) posterior.

**Figure 3.**
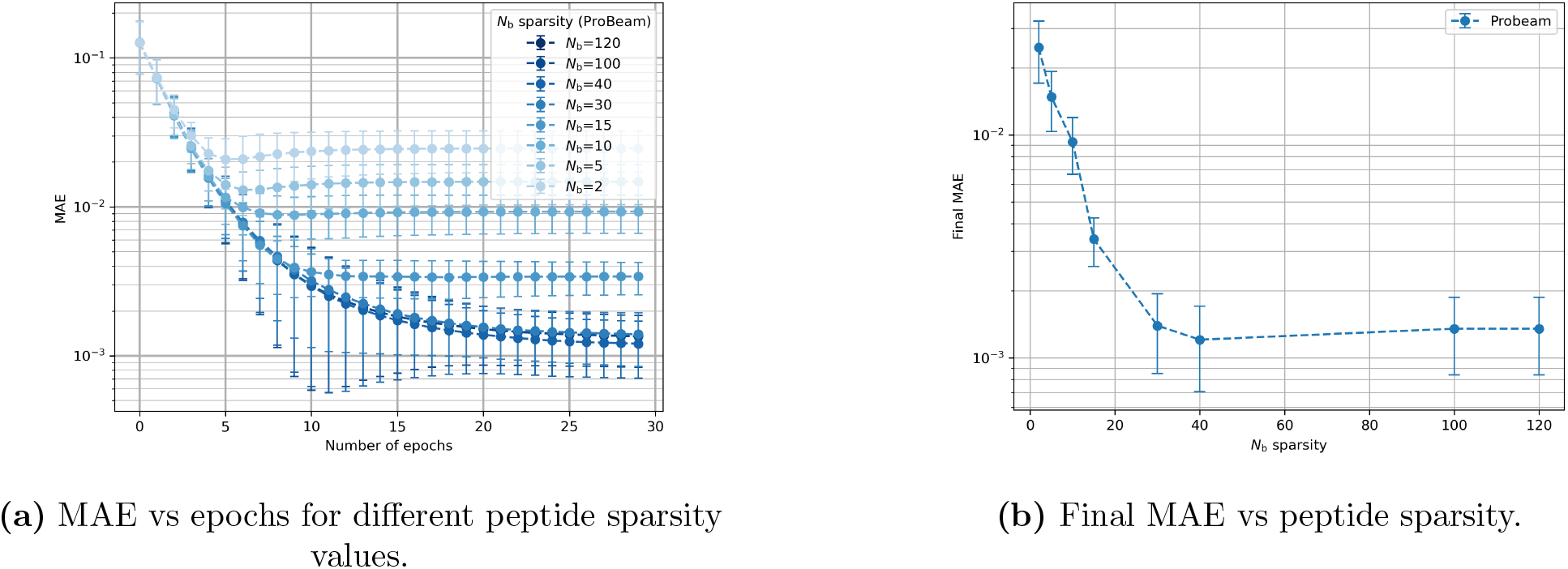
Effects of sparsity on convergence and final performance using Probeam for a 5-protein dataset.

### 4.3 Whole proteome protein inference

This section presents an evaluation of our algorithm’s ability to estimate protein abundances at the whole-proteome scale. The dataset was constructed from the human proteome as defined in UniProt, excluding only 18 proteins that did not generate any observable tryptic peptides, resulting in *N*_P_ = 20642 proteins.

#### 4.3.1 Comparison of abundance estimation

We now compare protein inference results obtained using the oracle and sparse Probeam (*N*_b_ = 1000), under both typical and reduced experimental error conditions and 10M reads. The sparsity value was chosen so that the computation times remained tractable and the convergence characteristic was illustrated correspondingly.

Figure 4 shows the convergence behavior of the expectation maximization algorithm for each case, with runtimes reported as the average per dataset.

**Figure 4.**
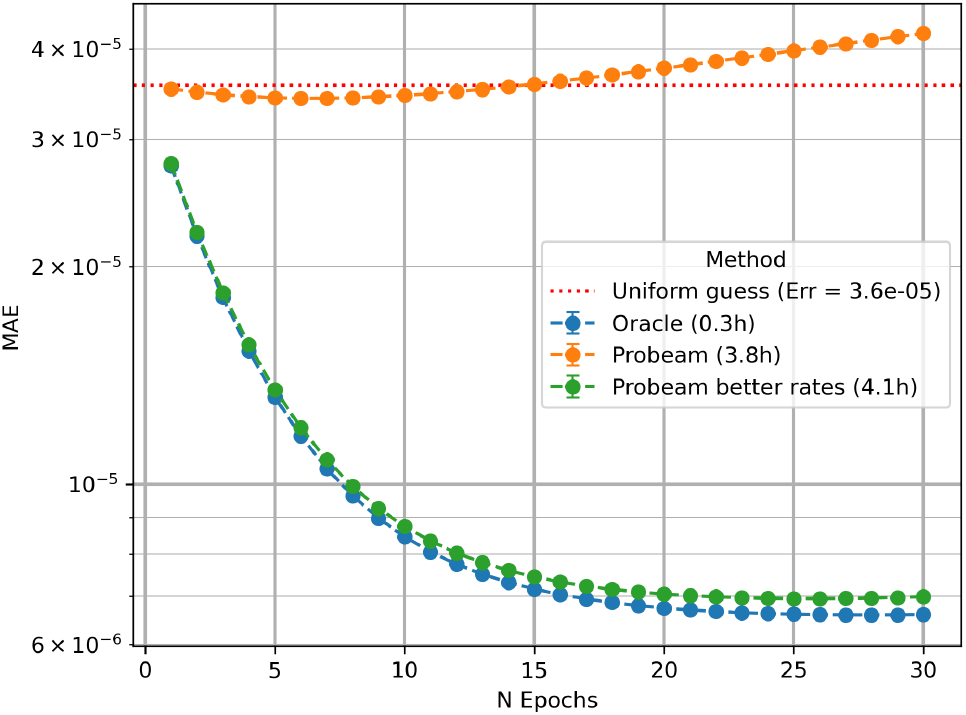
Convergence of EM algorithm for *N*_P_ = 20642, *N*_r_ = 10^7^.

Using the method described in Appendix 9.5, we estimate that each of the ten datasets contains approximately 3 *×* 10^5^ protein molecules using the method described in Appendix 9.5. This method also allows us to estimate the expected number of reads contributed by each protein, and we verified that, on average, 99.9% of proteins were represented by at least one read in each dataset.

The oracle achieves the lowest estimation error, establishing a best-case reference for inference performance. Under standard experimental conditions, Probeam initially performs better than random guessing; however, its estimation error increases with successive EM iterations. This degradation is consistent with the behavior observed in high-error oracle classifiers and can be attributed to the posterior estimates of Probeam not being sparse enough (see Appendix 9.4).

When experimental error rates are artificially reduced, Probeam produces substantially more accurate estimates. In this regime, the EM algorithm converges to protein abundance distributions that more closely reflect the ground truth. The simulation parameters used to evaluate this improved performance are provided in the Appendix 9.3.2. The oracle results can also be interpreted as the limiting case of Probeam as experimental error rates approach zero.

Finally, Figure 4 also highlights that oracle-based inference requires significantly less computational time than Probeam. Nevertheless, our EM algorithm remains computationally efficient and can be executed at whole-proteome scale using standard GPU resources.

Figure 5 visualizes the inferred protein distribution vs the true distribution for a single dataset:

**Figure 5.**
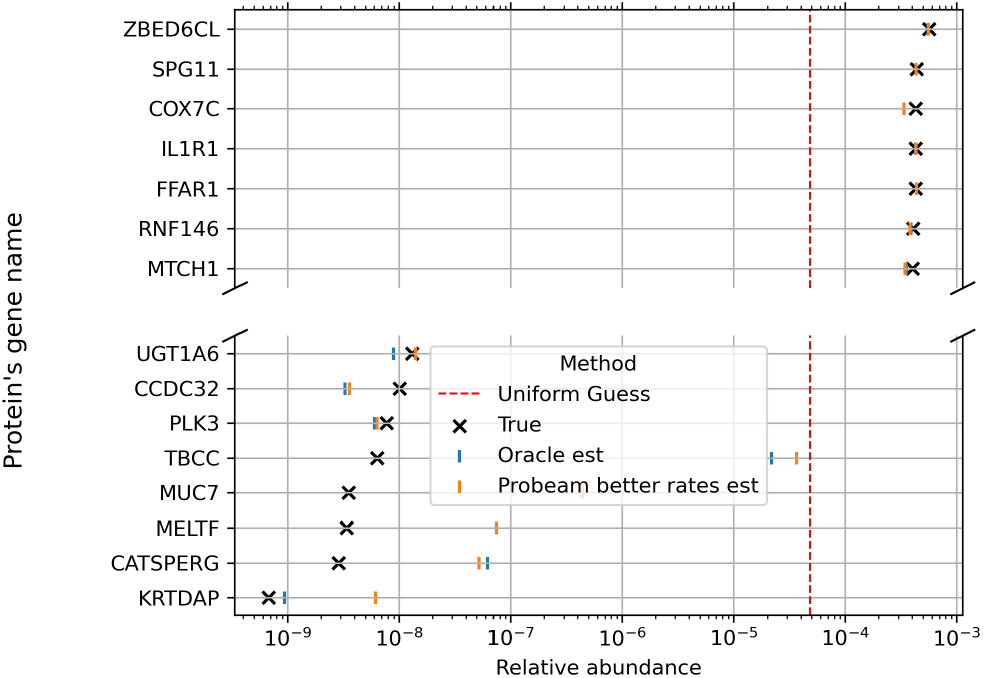
Example of a protein distribution estimation for *N*_P_ = 20642, *N*_r_ = 10^7^.

In Figure 5, proteins are sorted by their true abundances (*y*-axis), while the *x*-axis displays the true abundance values and the corresponding EM-based estimates after thirty epochs. The results show clearly that all classifiers outperform random guessing, with the oracle providing the closest match to the ground truth. For visualization clarity, only the improved-error version of Probeam is shown.

Although relative abundance estimates are less accurate in this large-scale setting than in the five-protein case, the algorithm still provides meaningful improvement over baseline methods. Importantly, our EM framework enhances inference for any estimation of the abundance. Therefore, it can be combined with other methods to achieve a more accurate protein inference on whole-proteome scale.

#### 4.3.2 Oracle performance

We analyze also the performance of the oracle classifier in this large-scale setting. Figure 6b shows how the final MAE varies with the number of reads, assuming a perfect peptide sequencing. Figure 6a illustrates the impact of different oracle’s classification error probabilities *e* on protein abundance estimation.

**Figure 6.**
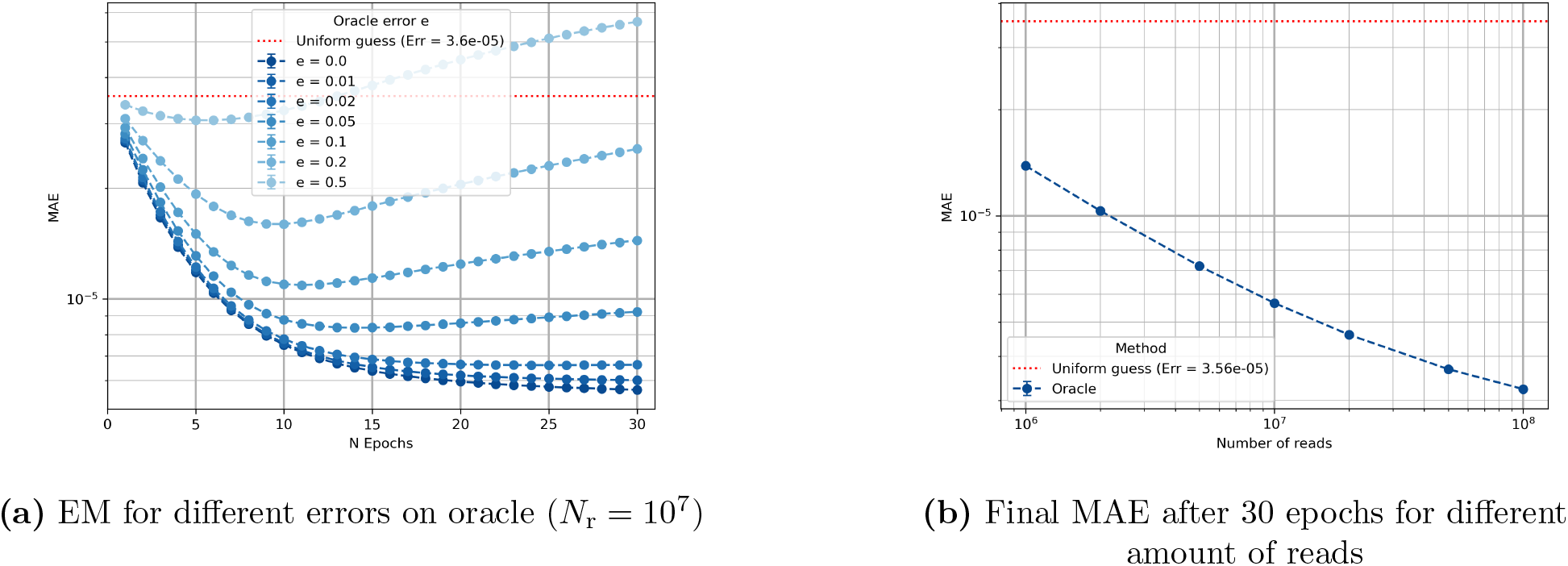
Sensitivity of the algorithm with respect to the number of reads and the classifier error rate.

As observed in the five-protein case, increasing the number of reads improves the quality of the abundance estimates. Similarly, increasing the classification error degrades performance, with convergence curves again showing non-monotonic behavior. One key difference in this full-proteome setting is that the error bars in the plots are negligible and not visually apparent, indicating low variance across different datasets.

Overall, the perfect oracle represents an idealized case in which peptide classification is error-free. As expected, it leads to significantly more accurate abundance estimation and serves as a reference point for evaluating also protease and labelling schemes.

## 5 Discussion

In this work, we presented a protein inference framework based on expectation maximization (EM), specifically designed for fluorosequencing data. Through simulations, we demonstrated that the method robustly estimates protein abundances under realistic experimental error rates, particularly in small-scale protein settings. Moreover, we showed that the algorithm is scalable to full-proteome datasets, making it compatible with the demands of fluorosequencing applications.

This scalability is especially important given the goals of fluorosequencing, which aims to perform single-molecule protein sequencing with ultra-high throughput, a level of performance that traditional proteomics methods cannot achieve. Efficient and scalable inference algorithms, like the EM-based approach proposed here, are essential to unlock the full potential of this technology in real-world applications. Furthermore, this framework could serve as a basis for other similar technologies using peptide-level information through nanopores [6, 18].

Despite these promising results, there is still room for improvement both in terms of estimation accuracy and computational efficiency, particularly for large protein sets.

From an accuracy standpoint, future work could explore the integration of expert knowledge into the prior distribution over proteins. Incorporating such priors could help improve the inference and guide it in biologically plausible directions. In addition, this EM-based method could serve as a refinement step for other inference pipelines, improving their final estimates. Another avenue would be to devise efficient approximations of the full data likelihood. Such computations would make it feasible to run EM from multiple initial protein distributions and, after convergence, rank the resulting solutions by their likelihood, retaining the one that achieves the highest local maximum.

On the performance side, additional sparsification strategies could be investigated to further reduce memory usage and computational load during EM iterations. More broadly, research into EM acceleration techniques -such as variational EM, stochastic EM, or quasi-Newton updates-could yield substantial improvements in convergence speed. Additionally, incorporating partial updates of the abundance estimates between batches could further accelerate convergence, since the GPU implementation processes batches of reads sequentially and updating the estimates is computationally lightweight.

In this study, all simulations were performed using a fixed experimental setup that included a specific choice of labeled amino acids, a single protease, and three distinct fluorescent dye channels. However, alternative experimental configurations, such as increasing the number of fluorophores or selecting different amino acids for labeling, could enhance the resolution of peptide-level data. For instance, adding a fourth dye would potentially improve the information of observable fluorescence patterns, which could improve peptide discrimination and, by extension, the accuracy of protein abundance estimates. Exploring such variations in experimental design represents a promising direction for future work, particularly in conjunction with the proposed inference framework.

Finally, the integration of deep learning into this pipeline presents a compelling opportunity. Neural networks could be trained to predict protein abundances directly from the raw fluorescence reads, bypassing some of the intermediate steps entirely. Alternatively, one could learn the *η* values in Equation 13 used in the EM updates using a neural network, allowing the EM framework to function in conjunction with learned components. Such hybrid or full data-driven approaches may not only accelerate inference but also increase robustness to modeling mismatches and experimental noise, effectively adapting the model to the data.

## 6 Method

### 6.1 Fluorosequencing

Fluorosequencing is a single-molecule protein sequencing (SMPS) technique inspired by methods used in DNA and RNA sequencing [9][10]. The process begins by denaturing proteins and proteolytically cleaving them into peptides, which are then chemically labeled with fluorescent dyes. Millions of these labeled peptides are immobilized in a flow cell and imaged using total internal reflection fluorescence (TIRF) microscopy.

Edman degradation is then performed in cycles, sequentially removing one N-terminal amino acid from each peptide. After each cycle, the fluorescence intensities of the peptides are measured. The pattern of fluorescence intensity drops across cycles is then analyzed by a peptide inference algorithm, which estimates the likelihoods of possible peptides generating that pattern. The possible peptides are known because the proteins in the sample, the labeling chemistry, and the proteolysis rules governing peptide cleavage are all defined.

It is important to emphasize that several systematic sources of error must be considered during the peptide inference step. These include: failure of dyes to attach to the targeted amino acids, failure of the Edman degradation process, detachment of peptides from the substrate, loss of dyes following degradation cycles, measurement noise in the fluorescence signal, and a blocking effect, a phenomenon in which a disruption prevents further Edman degradation from proceeding. Each of these factors introduces uncertainty into the observed fluorescence sequences and must be taken into account in realistic modeling and downstream inference.

An additional complication in fluorosequencing is that not all peptides are experimentally observable. This non-observability arises from several factors. First, some peptides may lack any of the specific amino acids targeted for dye labeling, making them inherently undetectable by the optical system. Even when labelable residues are present, there remains a non-negligible probability that all corresponding dye attachment reactions fail, resulting in fluorescence sequences with no detectable signal. Furthermore, peptides must be successfully immobilized on the surface of the flow cell at an appropriate distance from neighboring peptides; some may fail to bind altogether, while others may bind too close to adjacent molecules and be excluded during image processing.

Once peptide sequences and their probabilities are inferred from the observed fluorescence reads, they can be mapped back to their corresponding proteins, allowing for the estimation of protein abundances in a sample. This inference step also relies on a known database of proteins, which provides the expected peptide-protein relationships and helps resolve ambiguities in peptide assignments.

### 6.2 Protein abundance inference in Fluorosequencing

The random variable ***X*** represents the whole dataset generated by fluorosequencing a sample, which consists of *N*_r_ different reads. Each read, *X*_*k*_, is a matrix that contains the recorded light intensity after each Edman degradation sample for different colors. The rows of *X*_*k*_ correspond to the different fluorophore channels (colors), while the columns represent the sequential cycles of Edman degradation performed on the peptide. We define ***x*** as an observation of the whole dataset, while *x*_*k*_ is an observation of the *k*th read.

The reads *X*_*k*_ are generated by different peptides attached to the wall. Certain peptides are mutually indistinguishable by fluorosequencing, and the indistinguishable groups are referred to as fluorescence strings. The number of possible fluorescence strings is finite and is determined by several factors: the proteins utilized, the enzymes employed to digest the proteins (e.g., trypsin), and the specific amino acids labeled with fluorophores. We introduce the random variable *F* , representing a uniformly drawn fluorescence string in the experiment, with the domain of all possible fluorescence strings denoted as 𝒟_F_, with | 𝒟_F_| different fluorescence strings. The distribution of *F* is given by *P*_*F*_ (*f*). Note that some peptides may lack the specific amino acids required for fluorophore attachment. As a result, these peptides are represented by a fluorescence string that contains no dyes, referred to as the null fluorescence string (*f*_null_).

The purpose of protein abundance inference is to be able to estimate the protein distribution present in a solution. Let *Y* ∈ 𝒟_Y_ be a random variable representing a uniformly drawn protein, and *P*_*Y*_ (*y*) the distribution of proteins, which is the target for estimation. Given that the number of possible proteins, *N*_P_, is known for our experiments, we can represent *P*_*Y*_ (*y*) as follows:

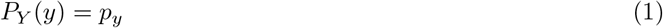

where *p*_*y*_ represents the relative abundance of the *y*th protein, where Σ _*y*_ *p*_*y*_ = 1.

Given a known protein distribution, the fluorescence string distribution can be inferred a priori. Specifically, since we know how each protein is cleaved and labeled, we can determine the relative frequency of fluorescence strings generated by each protein and extend this process to the entire protein set.

The protein inference problem can be formulated as estimating the protein distribution that maximizes the likelihood of the observed dataset:

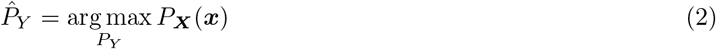

Solving Equation 2 is a highly complex problem, and in this paper, we present a method for obtaining practical estimates of 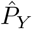. While not necessarily optimal, the proposed method provides meaningful estimations and can be also used to refine other inference techniques.

### 6.3 Notation

To introduce our protein inference method, which is based on expectation maximization (EM), we first define additional auxiliary variables. A challenge in applying EM for protein inference in fluorosequencing is that the number of proteins does not directly correspond to the number of reads: each protein can generate a different number of fluorescence strings, and some of these fluorescence strings may not be observable. This mismatch is a problem since the classic EM assumes the same number of hidden and observed variables, a problem that will be solved with our framework.

We begin by introducing the random variable *F* ^o^, which represents a fluorescence string that is experimentally observable. This variable can take the same values as *F* (the general fluorescence string variable), so *F* ^o^ ∈ 𝒟_F_. We consider 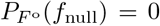 where *f*_null_ is the fluorescence string that represents no dyes on the string. The probability of dyes not attaching and the difference with *f*_null_ result in 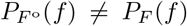, and an example is provided in Appendix 9.1 to further clarify the variable *F* ^o^. However, we can compute 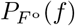 from a transformation of *P*_*F*_ (*f*):

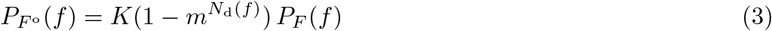

where *m* is the dye miss rate, *N*_d_ is a function that returns the number of amino acids that are ideally labeled and *K* is a normalization constant ensuring that the result is a valid probability 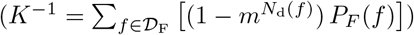.

Notice that *N*_d_(*f*_null_) = 0, and then the probability of observing the null fluorescence string is zero too. The advantage of *F* ^o^ is that there is one realization for each read in the dataset, and that 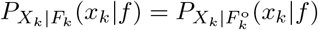 as the fluorescence string itself is unchanged. Finally, we define 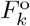 as the experimentally observed fluorescence string that generated the *k*th read.

Next, we introduce the protein indicator random variable *I*, which represents the original protein that generated a given read. The probability mass function *P*_*I*_ (*y*) describes the a priori likelihood that a read originates from protein *y*. For clarity, we denote *I*_*k*_ as the protein indicator for the *k*th read. An example is provided in Appendix 9.1 to further illustrate the concept of *I*. Both *I*_*k*_ and 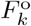 are hidden variables, while *X*_*k*_ is observable. The relationship between these variables is illustrated in Figure 7.

**Figure 7.**
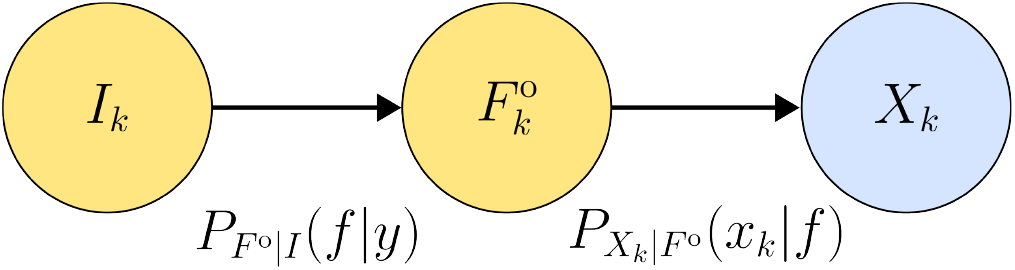
Relationship between the hidden protein Indicator *I*_*k*_, the hidden experimentally observable fluorescence string 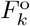 and the observable read *X*_*k*_ for the *k*th read.

The protein indicator mass probability function *P*_*I*_ can be described as follows:

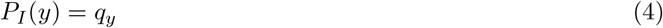

Each probability *q*_*y*_ is related to the original protein distribution *P*_*Y*_ (*y*) as:

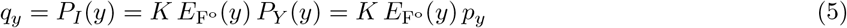

where 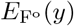 returns the average number of experimentally observable fluorescence strings generated by the protein *y*, and *K* is a normalizing factor defined as 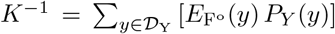. Equation 5 shows that knowledge of *P*_*Y*_ uniquely determines *P*_*I*_ , and also knowing *P*_*I*_ allows for the reconstruction of *P*_*Y*_.

Next, the distribution of experimental fluorescence strings for a given set of protein abundances can be expressed using the above introduced terms. This formulation is essential for generating the datasets used in testing:

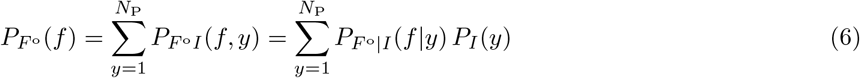

where *P*_*I*_ (*y*) is obtained from the protein distribution with Equation 5 and 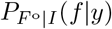 is known. Finally, we define 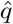 the estimated probabilities of the distribution of protein indicators:

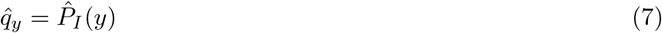

In summary, the protein indicator *I* is a hidden random variable with a realization for each observed read. This setup allows us to formulate an EM algorithm to maximize the likelihood of the observed data and estimate 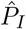. After fitting, the final step is to apply the inverse transformation from Equation 5 to obtain the estimated 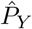.

### 6.4 Expectation maximization iteration

The Expectation-Maximization (EM) algorithm is an iterative technique used to find local maximum likelihood estimates of parameters in statistical models. It has numerous applications, including protein inference methods with mass spectrometry. In this paper, we apply the EM algorithm to iteratively update the estimate of the protein distribution, 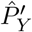, from a previous estimate, 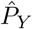, such that

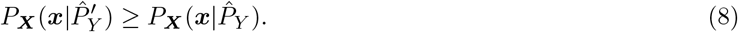

This iteration is derived by first noting that 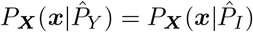 since the relation a known *P*_*Y*_ uniquely determines *P*_*I*_ with Equation 5. Consequently, the EM iteration can be formulated as:

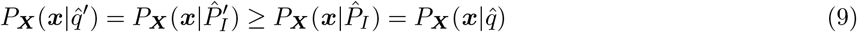

where 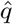 represents the parameters of the current estimated distribution 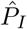 and 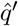 represents the parameters of the updated estimate 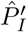. The inequality in Equation 9 is guaranteed by our algorithm, and the proof is provided in Appendix 9.6.

The parameter update equation is derived in Appendix 9.2, and shown in Equation 10:

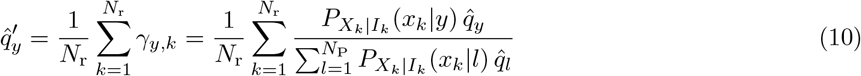

This result can be interpreted intuitively: *γ*_*y,k*_ can be seen as a joint probability between *X*_*k*_ and *I*_*k*_ when assuming a prior 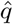 on *I*_*k*_, and then averaging over the reads results into a new estimate on the protein indicator distribution 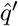.

The probabilities 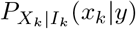 can be computed using the intermediate random variable 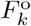 as follows:

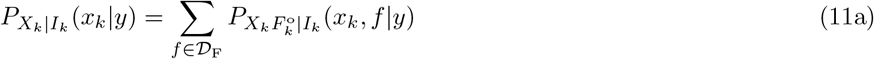

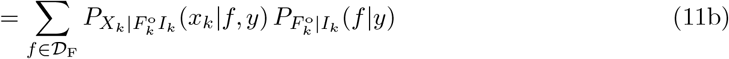

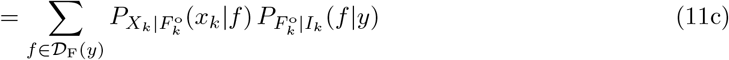

In Equation 11a, the expression is equal to the joint distribution of the reads with the experimental fluorescence strings marginalized over the experimental fluorescence strings. Next, we apply the chain rule in Equation 11b. Lastly we drop the conditioning on the indicator for the first probability because of the conditional independence illustrated in Figure 7 and we reduce the sum domain the possible fluorescence strings that can be generated by a protein *y*.

In the last step there are two probabilities inside the sum. First the reads likelihood is determined by the fluorescence string itself, as shown in Figure 7. Secondly the domain of the fluorescence strings can be reduced as the domain of fluorescence strings for a given protein *y*. This domain reduction significantly reduces the computational complexity, as each protein generates only a small subset of fluorescence strings.

### 6.5 Using the fluorescence string classification for the EM iteration

Algorithms such as Whatprot and Probeam allow us to estimate the distribution of fluorescence strings for a given read, but we analyze classifiers in general that can provide posterior estimates. For any classifier that assumes equally distributed fluorescence strings 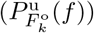 and provides a posterior estimate 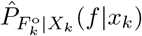, we can express

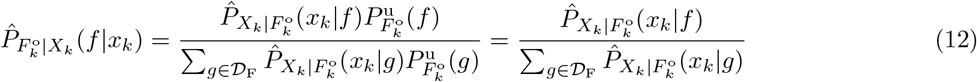

where the denominator can be written as a normalizing constant *C*_*k*_. Therefore 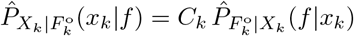. This result is important because it allows us to use the classificator’s posterior estimates to estimate 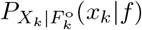 for the protein inference. The multiplicative term *C*_*k*_ is cancelled for the protein inference. Using this result and Equation 11 we can rewrite the EM update of Equation 10 as

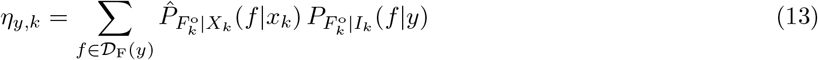

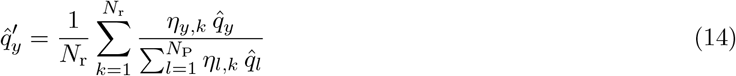

This equation shows how the updated protein distribution parameters 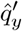 are obtained by summing over the reads and using the posterior probabilities from the classifier to update the EM steps.

### 6.6 Optimizing memory and processing time with sparsity on the fluorescence string classification

Computing the parameter updates for a large number of proteins, such as those in the human proteome, can demand significant computational resources. To address this, we leverage the sparsity of the posterior estimates 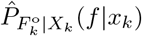. These estimates are sparse because, while there may be a large number of possible fluorescence strings (approximately 150K for the human proteome), only a small subset of fluorescence strings typically have a significant likelihood for any given read.

We model this sparsity as follows:

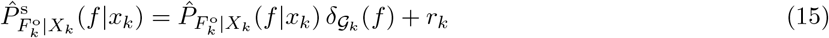

where the indicator function 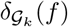 is defined as

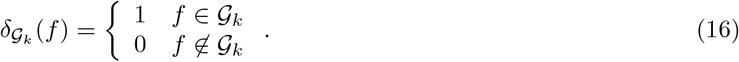

The set 𝒢_*k*_ contains the *N*_b_ fluorescence strings with the highest posterior probabilities for the read *k*; the indicator function selects only these top-ranked candidates for further computation. The indicator function selects therefore only the highest posterior probabilities for each read *k*.

The normalization constant *r*_*k*_ ensures that the remaining probability mass is uniformly distributed among all possible fluorescence strings, and is defined as 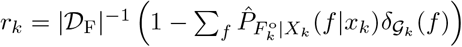. This sparsification strategy significantly reduces the computational and memory requirements for estimating *η*_*y,k*_.

Since we only need to consider fluorescence strings that are both produced by a given protein *y* (*f* ∈ 𝒟_F_(*y*)) and among the top *N*_b_ estimated posterior probabilities for a read *k* (𝒢_*k*_), we define the set 𝒟_F_(*y, k*) = {*f* ∈ (𝒟_F_(*y*) ∩ 𝒢_*k*_)*}* as the fluorescence strings that must be considered when approximating *η*_*y,k*_.

The sparsified form of the *η*_*y,k*_ computation becomes:

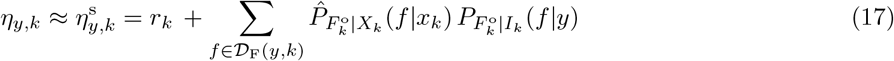

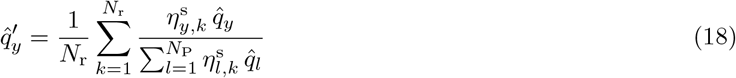

We evaluated various sparsification and normalization methods, as detailed in Appendix 9.7. This assessment was based on empirical tests, and the chosen method demonstrated the best performance. However, since the evaluation was conducted on small protein datasets, the results may vary for larger datasets. Nevertheless, this approach remains an optimized strategy that effectively reduces computational burden.

### 6.7 Posterior estimates

In this section it is described the different methods to obtain estimates of the posterior probabilities 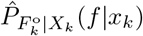

#### 6.7.1 Oracle

We define the oracle as an artificial posterior estimator designed for analytical benchmarking. This estimator has access to the ground-truth fluorescence string that generated each read in the simulation and allows us to characterize the upper performance bounds of the inference framework. The oracle posterior is defined as

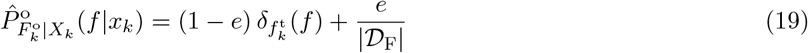

where 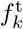 represents the true originating fluorescence string for the *k*th read, and *e* is the oracle’s error rate. This formulation ensures that the oracle assigns the highest probability to the true fluorescence string, while uniformly distributing the remaining probability mass across all other possible fluorescence sequences.

The oracle with zero error (*e* = 0) can be interpreted as an ideal fluorescence string inference method with perfect classification accuracy. This provides a useful theoretical baseline for evaluating the potential performance of the overall inference pipeline. For example, if the error-free oracle fails to yield accurate protein abundance estimates under certain experimental configurations (number of Edman degradation cycles, labeling strategies, and protease choices), then real-world posterior estimates, which inevitably introduce error, are unlikely to perform adequately under those conditions.

#### 6.7.2 Whatprot

Whatprot [14] is a two-stage classifier that first applies a k-Nearest Neighbors (k-NN) pre-selection to identify candidate fluorescence strings. Then, it computes the probability of the observed read given each candidate fluorescence string and, using Bayes’ rule, infers the posterior probability of each fluorescence string given the read.

While the Hidden Markov Model (HMM) in Whatprot allows for direct computation of 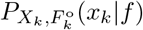, doing so for every possible fluorescence string remains computationally expensive. We used Whatprot to validate the accuracy of Probeam’s predictions and subsequently employed Probeam for the protein inference tasks.

Full posterior estimates over all fluorescence strings were required for our method, and modifying the Probeam implementation to output these estimates was more straightforward given our direct access to and familiarity with its code.

#### 6.7.3 Probeam

Probeam [15] was introduced as a computationally efficient alternative for fluorescence string classification, offering significant speed improvements at the cost of a modest reduction in accuracy. The method operates by defining a novel state-space and employing beam search decoding to approximate posterior probabilities over fluorescence strings.

However, Probeam was developed prior to the full characterization of the blocking effect, a sequencing artifact described in detail by [12]. In contrast, Whatprot now incorporates this phenomenon by adapting its Hidden Markov Models to account for its impact on fluorescence signal generation. As Probeam does not currently model blocking, we exclude this effect from our simulations. While this exclusion may lead to a slight overestimation of classification accuracy, its impact is minimal relative to the influence of other dominant error sources in the fluorosequencing process. While it is in principle possible to extend Probeam to account for blocking, implementing such enhancements is nontrivial and beyond the scope of the present work.

### 6.8 Error Metric

A standard approach for assessing parameter improvement in expectation maximization (EM) algorithms is to track the log-likelihood 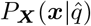 across successive iterations. In our setting, however, computing this likelihood is computationally intensive. As an alternative, we employ a distributional distance metric to evaluate the quality of the estimated protein abundance distribution.

Among the available options, we adopt the mean absolute error (MAE) for its simplicity, interpretability, and direct relevance to abundance estimation:

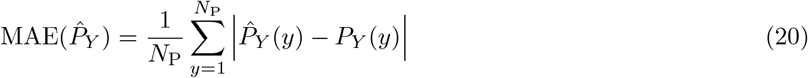

This metric quantifies the average deviation between the estimated and true protein distributions, providing an intuitive measure of estimation accuracy. While our EM algorithm guarantees non-decreasing likelihood with each iteration, it does not inherently ensure a monotonic improvement in MAE. Nonetheless, lower MAE values correspond to more accurate abundance estimates, making it a practical and informative proxy for performance evaluation in this context.

## 7 Acknowledgements

E.M.M. acknowledges grant support from the Welch Foundation (F-1515) and U.S. National Institutes of Health (R35 GM122480).

This work has also been supported by the Swedish Research Council (VR Research Environment Grant 2018-06169; QuantumSense), and the Swedish Foundation for Strategic Research (SSF Grant ITM17-0049).

ChatGPT was used to improve the cohesion, style, and vocabulary of the text, and Github Copilot was used for the development of the code.

## 9 Appendix

### 9.1 Illustrative examples of the presented random variables

#### 9.1.1 Experimental fluorophores

An example to illustrate the difference between *P*_*F*_ (*F* = *f*) and 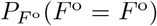 is the following. Suppose that for a setting of protein distribution, protein digestor and markers with fluorophore, we get *P*_*F*_ (*F* = *f*) as:

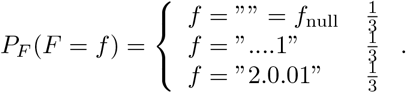

where ““ symbolizes the null fluorescence string. For example, the fluorescence string “2.0.01” represents a peptide that has two dyes of color 0, and one dye each of colors 1 and 2, arranged in the order of Edman degradation cycles, with the rightmost dye being the closest to the surface of the flow cell. This notation is described in more detail in prior work [14, 15].

The first step to obtain 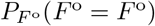 is to consider only the non-null dye sequences. We also have to consider that the dye sequences are not observable with a probability given by all the fluorophores not attaching. Since the dye miss probability was estimated in measurements to be *m* = 0.25, using Equation 3 we obtain that the distribution is:

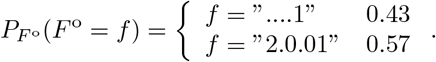

since *N*_d_(“….1”) = 1 and *N*_d_(“2.0.01”) = 4. Here we can observe that the first fluorescence string will appear less in the experiment because many times it wont have any fluorophore attached, while the second is more likely to have at least one fluorophore attached before the degradations. If they had the same amount of fluorophores, they would be equally likely.

#### 9.1.2 Protein indicator distributions

Here we show in table 1 an example to illustrate the difference between *P*_*Y*_ and *P*_*I*_:

**Table 1:** Protein indicator distribution example.

| Protein | $P_Y(Y = y)$ | $E_{F^\circ}(y)$ | $P_I(I = y)$ |
| --- | --- | --- | --- |
| 1 | 0.5 | 7.5 | 0.75 |
| 2 | 0.5 | 2.5 | 0.25 |

**Table 2:** Simulation parameters for the five-protein dataset.

| Parameter | Value |
| --- | --- |
| Number of Edman cycles | 11 |
| Edman failure probability | 0.06 |
| Detach probability | 0.05 |
| Initial blocking probability | 0 |
| Cyclic blocking probability | 0 |
| Dye bleach probability (all fluorophores) | 0.05 |
| Dud probability (all fluorophores) | 0.07 |
| Mean Gaussian light intensity | 10000 |
| Standard deviation of Gaussian intensity | 1600 |
| Standard deviation of background noise | 66.7 |

Note that the average number of experimental fluorescence strings for a protein can be not integer. In this example, protein 1 and 2 are equally distributed, but the first one produces three times more experimental fluorescence strings than the second. This effect leads to the given distribution of the protein indicator; it is more likely that an experimentally observable fluorescence string came from the first protein than from the second.

### 9.2 EM iterative step proof

In this subsection, we derive the parameter update equation for the Expectation-Maximization (EM) algorithm in fluorosequencing. The function 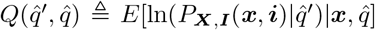 when maximized with respect to 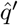 increases the likelihood of the reads given the new parameters 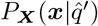, as demonstrated in Section 9.6. Here, we first rewrite the function 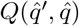, then maximize it, and finally derive the parameter update rule that results from this maximization.

To proceed, we define some useful notation. The realizations 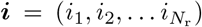 represents the protein indicators realizations for each read. We also introduce the vector 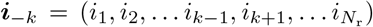 which excludes the indicator for a specific read *k*. With this notation, we rewrite the function *Q* as:

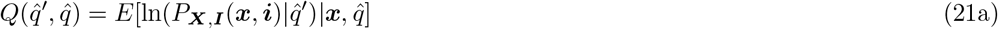

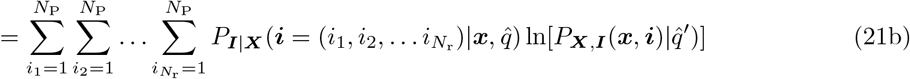

In Equation 21, we applied the expectation definition by summing over all possible values of the hidden protein indicator vector ***i***. Next, we expand ln 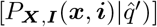:

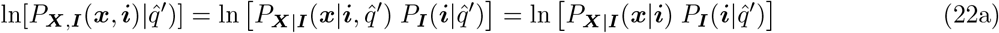

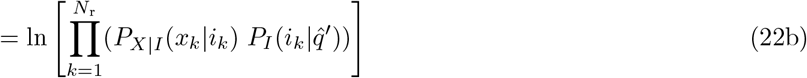

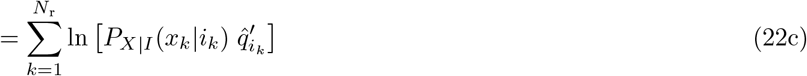

Equation 22a follows from the chain rule and the fact that ***X*** are observations conditioned only on the hidden variables ***I***. In Equation 22b we use the independency between the different reads and that we can drop the dependancy on the abundance estimations. Finally, in Equation 22c, we use the logarithm properties to make the product an external sum.

Consequently, substituting the result of Equation 22c in Equation 21b, we obtain:

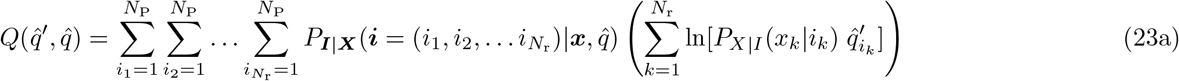

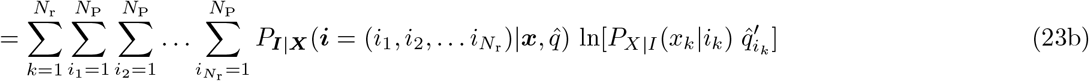

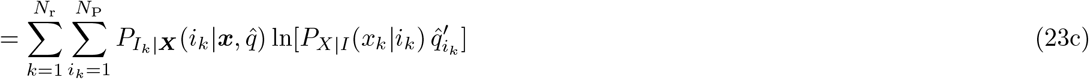

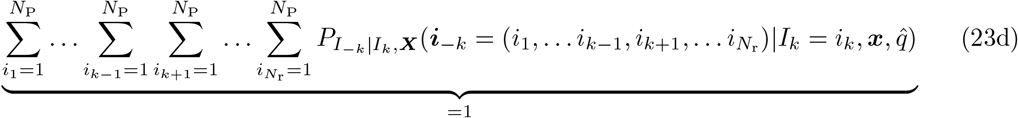

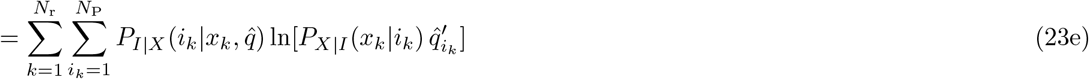

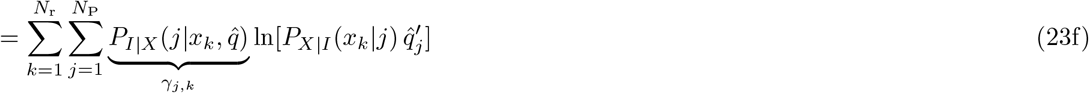

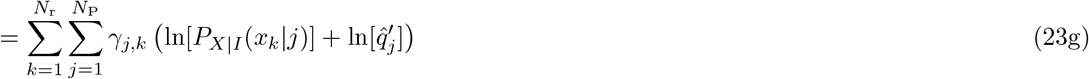

Here, in Equation 23b, we reorganize the summations. Equation 23c applies the chain rule for the indicator variable of read *k* (*i*_*k*_), and rearranges the terms. In Equation 23e the indicator is dependent only on the *k*th, so the condition can be dropped from ***X*** to *X*_*k*_. In Equation 23e, we exploit the fact that the indicator variable is independent of other reads, allowing us to drop the conditioning only to *x*_*k*_. Finally, in Equation 23g, we arrive at the final expression for 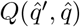, which serves as the basis for deriving the EM parameter update rule.

We simplify *γ*_*j,k*_:

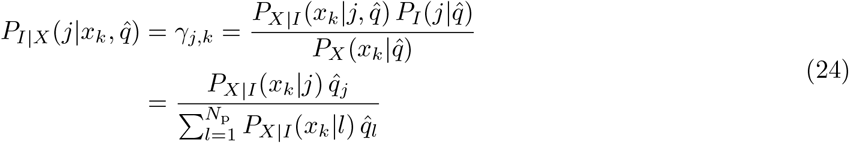

To maximize 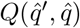 over 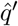, we need to introduce a constraint because the parameters 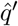 have to sum up to one. Therefore, using a Lagrange multiplier *λ*, we define the funtion 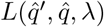 as:

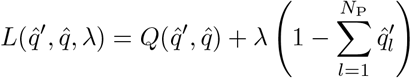

Taking the derivative with respect to 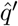 and setting it to zero, we obtain:

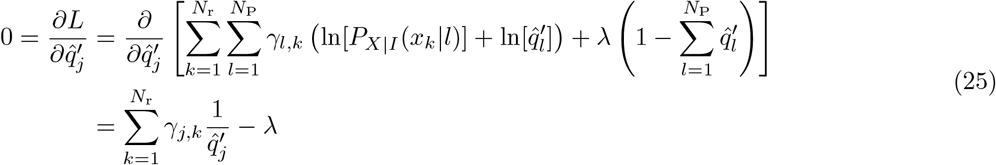

Rearranging gives:

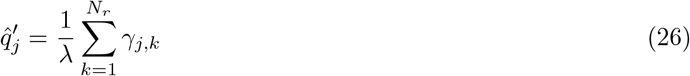

Since the sum of all 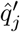 must be 1, we solve for *λ* and find:

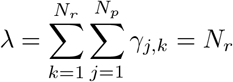

Equation 26 shows the final result on how to update the parameters in each EM epoch.

### 9.3 Simulation parameters

#### 9.3.1 Five proteins

The simulation parameters used for the five-protein experiments are consistent with those reported in [14] and [15], with one exception: the number of Edman degradation cycles was set to 11. This adjustment ensures that each fluorescence string is distinguishable by the sequencing platform.

We note that blocking probabilities (initial and cyclic) were not included in these simulations, as they are not implemented in Probeam. Nevertheless, their exclusion does not qualitatively affect the results.

#### 9.3.2 Whole proteome

For the whole-proteome experiments, we simulated fluorescence reads under two distinct experimental configurations. The first reflects typical error rates observed in current fluorosequencing platforms, consistent with those used by Probeam. The second adopts significantly reduced error rates to assess inference performance under near-ideal experimental conditions. All the error rates are shown in Table 3.

**Table 3:** Simulation parameters for the whole-proteome dataset under standard and improved error conditions.

| Parameter | Standard Error (Probeam) | Improved Error |
| --- | --- | --- |
| Number of Edman cycles | 39 | 39 |
| Edman failure probability | 0.06 | 0.0006 |
| Detach probability | 0.05 | 0.0005 |
| Initial blocking probability | 0 | 0 |
| Cyclic blocking probability | 0 | 0 |
| Dye bleach probability (all fluorophores) | 0.05 | 0.0005 |
| Dud probability (all fluorophores) | 0.07 | 0.0007 |
| Mean Gaussian light intensity | 10000 | 10000 |
| Standard deviation of Gaussian intensity | 1600 | 160 |
| Standard deviation of background noise | 66.7 | 66.7 |

The number of Edman degradation cycles was set to 39 in both configurations. Shorter sequences led to substantial overlap among fluorescence strings, making many of them indistinguishable and thereby drastically reducing inference accuracy.

### 9.4 Probeam estimates: additional information

In this section, we compare the classification accuracy of Probeam and Whatprot to justify the use of Probeam as a surrogate for Whatprot in our experiments. Probeam was always run with 90 beams. As shown in Table 4, both methods exhibit similar accuracy across different datasets and error configurations. In some cases, Probeam even marginally outperforms Whatprot.

**Table 4:** Comparison of Probeam and Whatprot classification accuracy across datasets and error configurations.

| Number of Proteins | Error Parameters | Probeam Accuracy | Whatprot Accuracy |
| --- | --- | --- | --- |
| 5 | Standard | 61.93% | 61.94% |
| 20,642 | Standard | 9.85% | 9.83% |
| 20,642 | Improved | 98.19% | 98.15% |

To observe the sparsity the posterior estimates, we accumulate the posterior estimates ordered by their probability, and we obtain the average residue error for each *N*_b_ of sparsity. This is done for the three different Probeam we use and it is shown in Figures 8, 9 and 10.

**Figure 8.**
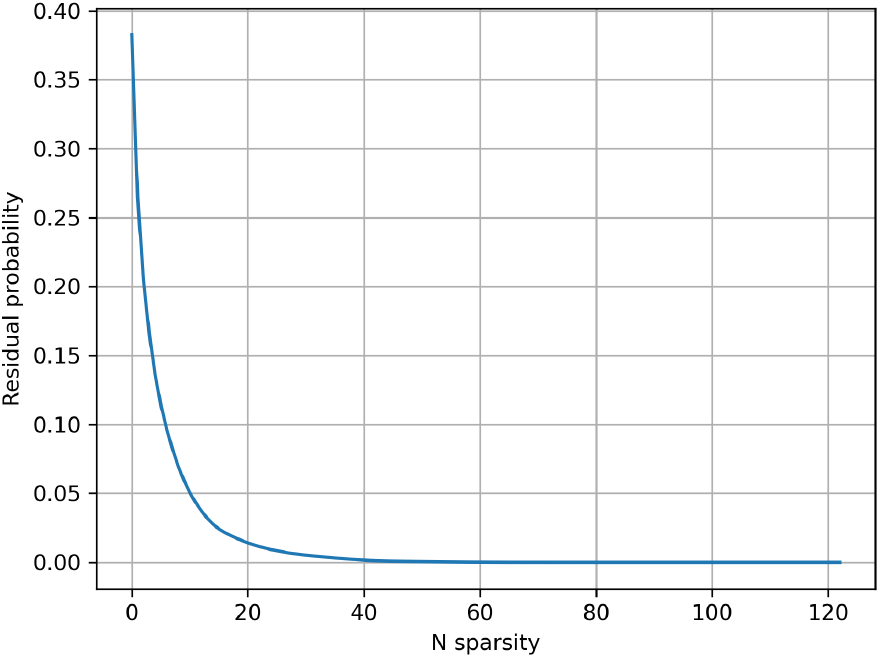
Average residue probability for sparse Probeam posterior estimates (standard error rates) on the five-protein dataset.

**Figure 9.**
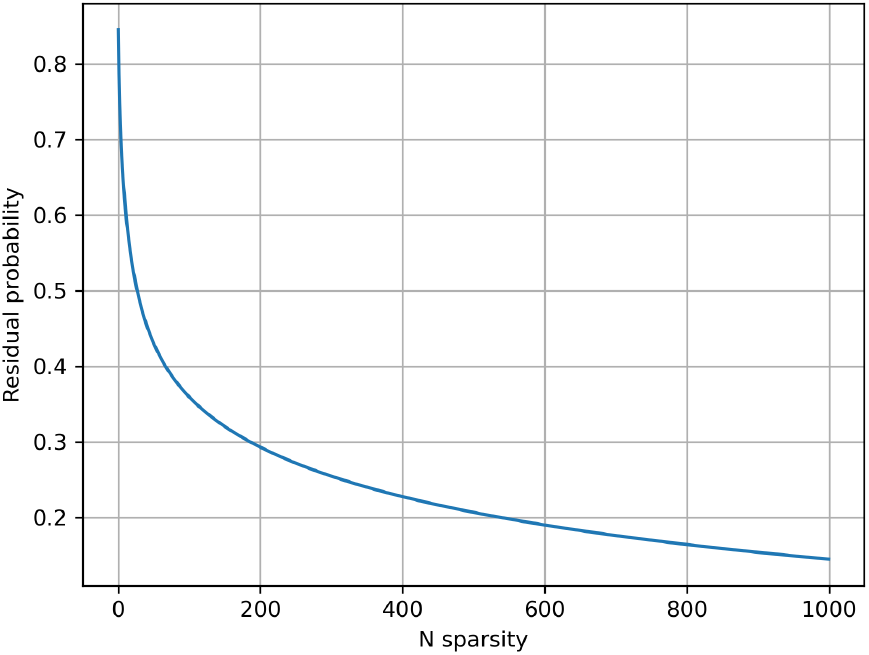
Average residue probability for sparse Probeam posterior estimates (standard error rates) on the whole proteome.

**Figure 10.**
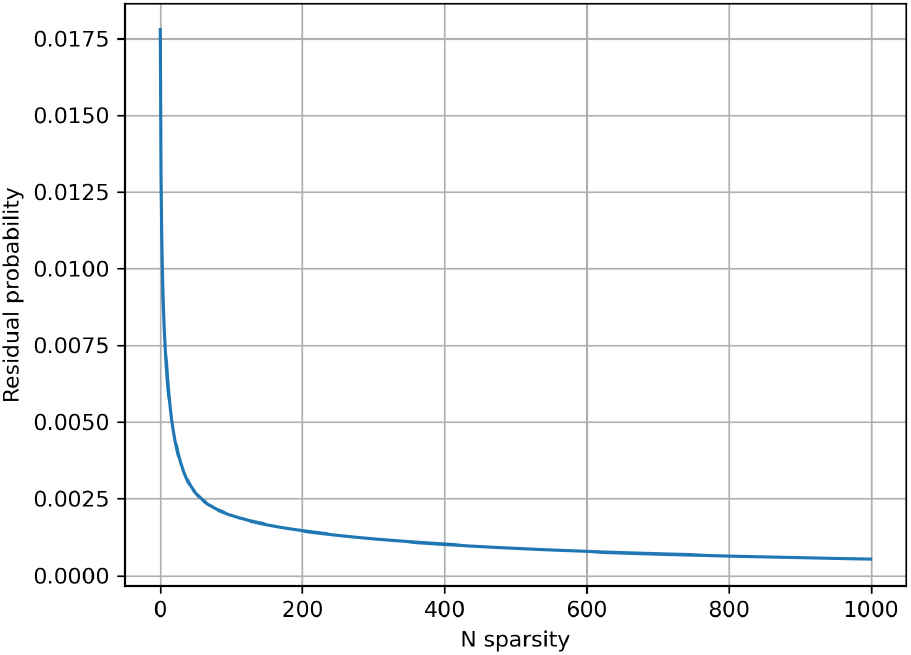
Average residue probability for sparse Probeam posterior estimates (improved error rates) on the whole proteome.

To better understand the sparsity of the posterior distributions generated by Probeam, we evaluate the average residue probability as a function of the number of top *N*_b_ fluorescence strings retained. The *x*-axis represents the sparsity value, and *y*-axis is the average residual probability for the whole shared dataset. This analysis is presented in Figures 8, 9, and 10 for the three Probeam configurations used.

Figure 8 shows that the posterior estimates for the five-protein dataset are highly sparse, which explains why a small *N*_b_ still provides a good approximation. In contrast, Figure 9 reveals that even after retaining the top 1000 posterior entries, the average residue remains above 0.1. This indicates that the posterior is less concentrated, which contributes to the poorer inference performance under standard error conditions on the whole proteome. On the other hand, Figure 10 shows much lower residue values, confirming that the improved error rates result in more peaked and informative posterior distributions.

### 9.5 Estimating protein counts

The total number of proteins in the sample, denoted *K*_P_, can be estimated using the following expression:

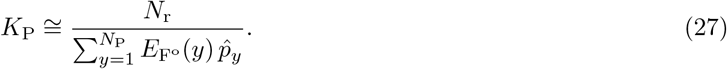

where *N*_r_ is the total number of reads, 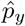 is the estimated relative abundance of protein *y*, and 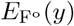 is the expected number of observable fluorescence strings generated by protein *y*.

Since we do not observe which protein generated each individual read, we approximate that each protein contributes its expected number of observable reads. The protein count for a particular protein *y* is given by 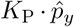, and the total number of reads observed is *N*_r_. By adding the expected contributions across all proteins and rearranging the terms, we arrive at Equation 27.

Applying Equation 27 to our human proteome datasets with *N*_r_ = 10^7^, we obtain an average estimate of *K*_P_ *≈* 3 *×* 10^5^.

### 9.6 Maximization proof

This proof demonstrates a key property of the Expectation-Maximization (EM) algorithm: maximizing the auxiliary function 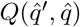 over 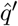 guarantees an increase in the likelihood of the observed data under the new parameters 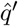. Our proof adapts arguments from [19] to the context of fluorosequencing. Specifically, we aim to demonstrate that:

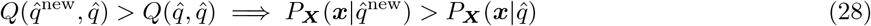

where 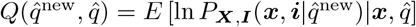, and ***i*** represents the hidden protein indicators.

We begin by comparing the log-likelihood of the data under two parameter sets 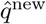 and 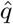:

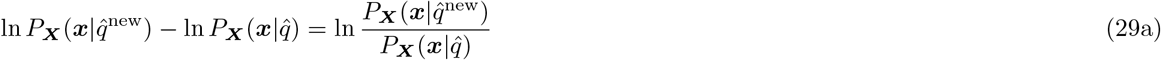

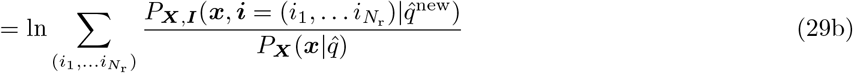

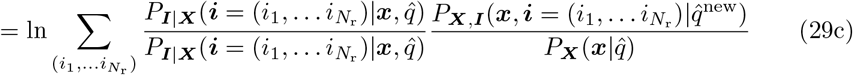

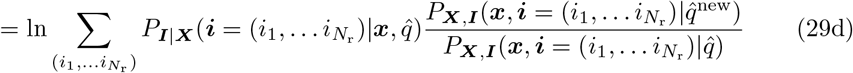

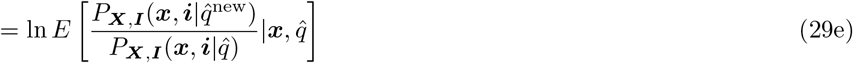

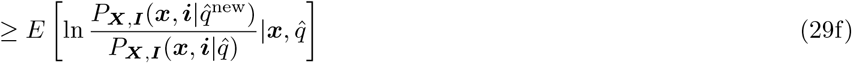

In Equation 29b, we marginalize over the hidden protein indicator variables. Then, in Equation 29c, we multiply and divide by the same term to facilitate a useful regrouping of variables. Moving to Equation 29e, we rewrite the summation as an expectation, which allows us to apply Jensen’s inequality in Equation 29f. Since the logarithm is a concave function, applying Jensen’s inequality ensures that the expectation of the logarithm is less than or equal to the logarithm of the expectation, leading to the inequality in the final step of this derivation.

Next, we rewrite the expectation obtained in Equation 29f in terms of the function *Q*:

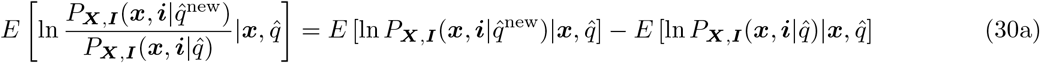

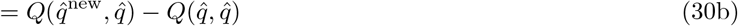

By combining Equation 29f with Equation 30b, we obtain:

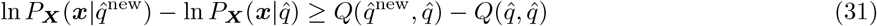

Equation 31 proves the original proposition stated on Equation 28.

### 9.7 Sparsity analysis

We first evaluated memory usage and performance across three different sparsification strategies:

- Retaining the top *N*_b_ most likely fluorescence strings.
- Retaining the most likely fluorescence strings whose cumulative likelihood remained below a specified threshold.
- Retaining all fluorescence strings with likelihoods above a predefined threshold.

Among these approaches, the top *N*_b_ most likely fluorescence strings method achieved the best balance, significantly reducing memory usage while maintaining negligible performance loss. In contrast, the other methods required significantly more memory to achieve a similar level of performance.

For normalization, we compared the following approaches:

- No normalization.
- Scaling scores to sum to one.
- Distributing the remaining probability uniformly across all fluorescence strings.
- Distributing the remaining probability uniformly across all fluorescence strings not included among the top *N*_b_ sparse entries.

The best results on the five-protein datasets were obtained using the third and fourth approaches. Distributing the remaining probability uniformly across all fluorescence strings (third approach) has the additional advantage of simplifying the residual term in Equation 17, allowing it to be factored out of the summation. The difference between the third and fourth approaches is minimal when the classifier output is already sparse and the number of possible fluorescence strings is large. For these reasons, we adopted the third approach in our experiments.

### 9.8 GPU Implementation comments

In our GPU implementation, we computed Equations 18 and 17. Since we process millions of reads simultaneously, it was infeasible to store all 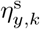 values directly; storing this matrix would require memory equal to *N*_r_ *× N*_P_ floats.

To address this limitation, we allocated as much GPU memory as possible to store the largest feasible subset of 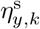. For the reads that fit in memory, we computed partial sums of Equation 18 in batches, accumulating these updates across all reads. After computing the whole epoch, the complete sum is obtained and the estimated abundances are updated.

When optimizing with CuPy, we found that the sparse matrix multiplication in Equation 17 became a major bottleneck. To overcome this, we implemented the operation directly at the CUDA level for greater efficiency. Most operations leveraged functions from the cuBLAS library; however, the sparse multiplication required a custom CUDA kernel.

After exploring different kernel designs, we settled on an approach where each block computed the sum for a subset of proteins and reads. The probabilities 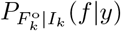 were loaded into shared memory, and each thread computed the sum for a corresponding read. We pre-sorted the indices of the sparse scores and the fluorescence strings associated with each protein. This preordering allowed faster index comparisons to identify overlapping indices between the protein and the top-scoring fluorescence strings. While the custom kernel achieved significant speed improvements, further optimization opportunities remain.

In the oracle case, where sparsity equals one, the matrix 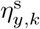 is also sparse. This greatly reduced memory access requirements during writes. By optimizing this specific case, we achieved runtimes comparable to cuBLAS routines. Therefore, further optimization of this case would yield only marginal additional speed gains.

## References

[1] Kübra Eren, Nursema Taktakoğlu, and Ibrahim Pirim. Dna sequencing methods: from past to present. The Eurasian journal of medicine, 54(Suppl 1):S47, 2022.

[2] W Timp and G Timp. Beyond mass spectrometry, the next step in proteomics. sci. adv. 6, eaax8978, 2020.

[3] Laura Restrepo-Pérez, Chirlmin Joo, and Cees Dekker. Paving the way to single-molecule protein sequencing. Nature nanotechnology, 13(9):786–796, 2018.

[4] Javier Antonio Alfaro, Peggy Bohländer, Mingjie Dai, Mike Filius, Cecil J Howard, Xander F Van Kooten, Shilo Ohayon, Adam Pomorski, Sonja Schmid, Aleksei Aksimentiev, et al. The emerging landscape of single-molecule protein sequencing technologies. Nature methods, 18(6):604–617, 2021.

[5] Luning Yu, Xinqi Kang, Fanjun Li, Behzad Mehrafrooz, Amr Makhamreh, Ali Fallahi, Joshua C Foster, Aleksei Aksimentiev, Min Chen, and Meni Wanunu. Unidirectional single-file transport of full-length proteins through a nanopore. Nature biotechnology, 41(8):1130–1139, 2023.

[6] Shengli Zhang, Gang Huang, Roderick Corstiaan Abraham Versloot, Bart Marlon Herwig Bruininks, Paulo Cesar Telles de Souza, Siewert-Jan Marrink, and Giovanni Maglia. Bottom-up fabrication of a proteasome– nanopore that unravels and processes single proteins. Nature chemistry, 13(12):1192–1199, 2021.

[7] Henry Brinkerhoff, Albert SW Kang, Jingqian Liu, Aleksei Aksimentiev, and Cees Dekker. Multiple rereads of single proteins at single–amino acid resolution using nanopores. Science, 374(6574):1509–1513, 2021.

[8] Brian D Reed, Michael J Meyer, Valentin Abramzon, Omer Ad, Omer Ad, Pat Adcock, Faisal R Ahmad, Gün Alppay, James A Ball, James Beach, et al. Real-time dynamic single-molecule protein sequencing on an integrated semiconductor device. Science, 378(6616):186–192, 2022.

[9] Jagannath Swaminathan, Alexander A Boulgakov, and Edward M Marcotte. A theoretical justification for single molecule peptide sequencing. PLoS computational biology, 11(2):e1004080, 2015.

[10] Jagannath Swaminathan, Alexander A Boulgakov, Erik T Hernandez, Angela M Bardo, James L Bachman, Joseph Marotta, Amber M Johnson, Eric V Anslyn, and Edward M Marcotte. Highly parallel single-molecule identification of proteins in zeptomole-scale mixtures. Nature biotechnology, 36(11):1076–1082, 2018.

[11] James H Mapes, Julia Stover, Heather D Stout, Tucker M Folsom, Emily Babcock, Sandra Loudwig, Christopher Martin, Mariah J Austin, Fan Tu, Casey J Howdieshell, et al. Robust and scalable single-molecule protein sequencing with fluorosequencing. bioRxiv, 2023.

[12] Matthew Beauregard Smith, Kent VanderVelden, Thomas Blom, Heather D Stout, James H Mapes, Tucker M Folsom, Christopher Martin, Angela M Bardo, and Edward M Marcotte. Estimating error rates for single molecule protein sequencing experiments. PLOS Computational Biology, 20(7):e1012258, 2024.

[13] James L Bachman, Christopher D Wight, Angela M Bardo, Amber M Johnson, Cyprian I Pavlich, Alexander J Boley, Holden R Wagner, Jagannath Swaminathan, Brent L Iverson, Edward M Marcotte, et al. Evaluating the effect of dye–dye interactions of xanthene-based fluorophores in the fluorosequencing of peptides. Bioconjugate chemistry, 33(6):1156–1165, 2022.

[14] Matthew Beauregard Smith, Zack Booth Simpson, and Edward M Marcotte. Amino acid sequence assignment from single molecule peptide sequencing data using a two-stage classifier. PLOS Computational Biology, 19(5):e1011157, 2023.

[15] Javier Kipen and Joakim Jaldén. Beam search decoder for enhancing sequence decoding speed in single-molecule peptide sequencing data. PLOS Computational Biology, 19(11):e1011345, 2023.

[16] Ting Huang, Jingjing Wang, Weichuan Yu, and Zengyou He. Protein inference: a review. Briefings in bioinformatics, 13(5):586–614, 2012.

[17] Enrique Audain, Julian Uszkoreit, Timo Sachsenberg, Julianus Pfeuffer, Xiao Liang, Henning Hermjakob, Aniel Sanchez, Martin Eisenacher, Knut Reinert, David L Tabb, et al. In-depth analysis of protein inference algorithms using multiple search engines and well-defined metrics. Journal of proteomics, 150:170–182, 2017.

[18] Henry Brinkerhoff, Albert SW Kang, Jingqian Liu, Aleksei Aksimentiev, and Cees Dekker. Multiple rereads of single proteins at single–amino acid resolution using nanopores. Science, 374(6574):1509–1513, 2021.

[19] Arne Leijon and Gustav Eje Henter. Pattern Recognition: Fundamental Theory and Exercise Problems. School of Electrical Engineering, KTH Royal Institute of Technology, Stockholm, Sweden, 2012. 2015 ed.

